# Visual Perception of 3D Space and Shape in Time - Part III 2D Shape Recognition by Log-Scaling

**DOI:** 10.1101/2022.03.01.482004

**Authors:** Brian Ta, Maria E. M. M. Silva, Kelly Bartlett, Umaima Afifa, Annie Agazaryan, Ricardo Canela, Javier Carmona, Emmanuel John L. De Leon, Alyssa Drost, Diego Espino, Guadalupe Espinoza, Kyleigh Follis, Paul Gan, Lauren Ho, Christina Honoré, Emily Huang, Luis Ibarra, Tessa Jackson, Mira Khosla, Caominh Le, Victor Li, Trevor McCarthy, Elizabeth Mills, Sukanya Mohapatra, Yuuki Morishige, Nancy Nguyen, Ziyan Peng, Kimya Peyvan, Michael Phipps, Isabella Poschl, Jagannathan Rangarajan, Charÿsa Santos, Leonard Schummer, Sky Shi, Natalie Smale, April Smith, Divya Sood, Cindy Ta, Anna Tran, Michelle Tran, Rui Wang, Patrick Wilson, Nicole L. Yang, Megan Yu, Selena Yu, Aaron P. Blaisdell, Katsushi Arisaka

## Abstract

Human vision has a remarkable ability to recognize complex 3D objects such as faces that appear with any size and 3D orientations at any 3D location. If we initially memorize a face only with a normalized size and viewed from directly head on, the direct comparison between the one-sized memory and a new incoming image would demand tremendous mental frame translations in 7D. How can we perform such a demanding task so promptly and reliably as we experience the objects in the world around us?

Intriguingly, our primary visual cortex exhibits a 2D retinotopy with a log-polar coordinate system, where scaling up/down of shape is converted to linear frame translation. As a result, mental scaling can be performed by linearly translating the memory or the perceptual image until they overlap with each other. According to our new model of **NHT** (Neural Holography Tomography), alpha brainwaves traveling at a constant speed can conduct this linear translation. With this scheme, every scaling up/down by a factor of two should take the same amount of extra mental time to recognize a smaller/larger face.

To test this hypothesis, we designed a reaction time (RT) experiment, where participants were first asked to memorize sets of unfamiliar faces with a given specific size (4° or 8°). Following the memorization phase, similar stimuli with a wide range of sizes (from 1° to 32°) were presented, and RTs were recorded. As predicted, the increase in RT was proportional to the scaling factor in the log scale. Furthermore, we observed that RTs were fastest for 8° faces even if the memorized face was 4°. This supports our hypothesis that we always memorize faces at the exact size of ~8 °. To our surprise, the increases in RT were also consistent with the mentally-estimated depth sensation, which indicates that the apparent size of the recognized face can create a proper depth sensation.

## 1 Introduction

We have a remarkable ability to quickly and effortlessly recognize a familiar face across a large range of distances from up close to far away. How we are able to do this has remained a mystery. The obvious first step is that we must memorize the face so that we may recognize it later. What is less well understood is how we can recognize a memorized face so well despite large changes in scale. The same question may be asked about the recognition of any familiar object (e.g., cars, trees, and buildings), and even written characters such as letters, numbers, and symbols.

What is remarkable is that the 2-dimensional (2D) retinotopic pattern created by an object, such as a face, in our visual field undergoes large distortions when the object is projected onto the retina from different distances. Recently, we proposed a theory that explains this capacity for rapid, efficient object recognition (see Arisaka, 2022a, b; Arisaka & Blaisdell, 2022). The starting principle of the theory is that the 2D spatial information contained in the retinotopic input becomes compressed to the temporal domain based on the principle of Neural Holographic Tomography (NHT). By encoding spatial information holographically in the time domain the many components of the observed face (or other objects) can be bound together in a single memory and stored in a holographic ring attractor lattice (HAL) structure in a compressed form. The compressed memory can then be expanded in the hippocampus where the current visual input can then be compared to the expanded memory. Thus, NHT and the HAL structure for memory provide the mechanisms by which top-down and bottom-up processes enable visual memory and recognition (Kinchla & Wolfe, 1979). The bottom-up process gathers sensory information from the environment while the top-down process begins with a memorized image of a previously encoded stimulus. The product of both processes are compared to identify an objects in our visual field.

Another benefit of NHT compression is that simple, linear processing of the HAL can mimic the scalar changes within the visual system that the perceived image undergoes when the real observed face (object) is viewed at various distances. That is, a log-polar scalar process can be automated in linear cartesian space so that perception and memory can be aligned to allow for rapid comparison and decision (e.g., “Yes, this is John”, or “No, I’ve never met her before”).

To achieve this scaling invariance, it has been found that a log-polar mapping exists for a retinal image projected to the cortex. Any transformation in scale or rotation causes the log-polar mapping in the cortical plane to shift 2-dimensionally (Araujo and Dias, 1996; Le et al., 2022). This aspect of vision is vital for pattern recognition and further implies that the perception of scaled images at varying heights are related according to the log scale. Once an object, such as a face, is memorized and stored holographically, recognizing the same face in the future involves a process of comparing the size-specific stored memory of the face (like a prototype), and comparing it to the currently perceived face. Such comparison requires that the observed face be the same size as the stored face. Scalar manipulation of the perceived face involves a transformation from space to time following the principles of NHT described above. This process utilizes alpha-phase procession whereby the 2D retinotopic image is scanned by alpha waves and converted into a holographic time code. Alpha waves travel within a narrow frequency band (~8-10 Hz), and therefore the larger the stimulus. The greater the difference between the observed image and the memorized template, the greater the distance the alpha waves must travel to scale up or down the perceived image, necessitating a longer time to complete the mapping process between observed and memorized face. This process makes the prediction that participant Reaction Times (RTs) will be fastest when the perceived image is the same size as the memorized one (no scaling necessary), while RTs should increase linearly as the perceived image increases or decreases in size. In other words, when we observe a smaller (or larger) face or object that is larger or smaller than the memorized face or object by a factor of two, this requires shifting the phase of the horizontally traveling alpha brainwave by 1 cm to the left (or right), so that the incoming face/object pattern and the memorized face/object pattern can overlap. We predict a direct relationship between processing time (and thus RT) and log of stimulus height will be observed. This effect has never been thoroughly assessed, especially for complex, facial stimuli or written characters, which is the goal of the current experiments.

We test the predictions of our holographic theory of visual object encoding and recognition using a behavioral psychophysics paradigm recording reaction time in human participants. In each experimental protocol, participants first memorized a set of three stimuli (e.g., 3 human faces, 3 familiar characters of the English alphabet, or 3 unfamiliar Chinese characters), where they had to press one of three buttons, each button corresponding to one of the three items in the set. During memorization, items were always presented at the same, reference size. Following the memorization phase, RTs were then collected for the same stimuli presented at a range of sizes from much smaller to much larger than the reference size. We predicted fastest RTs when the test image size was the same as the reference image size, while increases and decreases in test image size would produce larger RTs the greater is the size difference between reference and test image.

## 2 Results

### 2.1 Raw Data Analysis

To assess data quality, all raw participant data were first plotted jointly under Figure 1. All data were plotted using a log scaled x-axis with two linear fits applied, centering on either log_2_(8 deg) or log_2_(4 deg), depending on which size the stimuli were initially memorized as in the protocols. As participants differ in their base reaction times, a Cousineau normalization was applied to shift all participants’ fits towards the aggregate mean, without affecting the slopes. As seen in Figure 3, all participants across the 3 independent groups display similar slopes for each protocol, with clear banding into a V-shape pattern.

**Figure 1.**
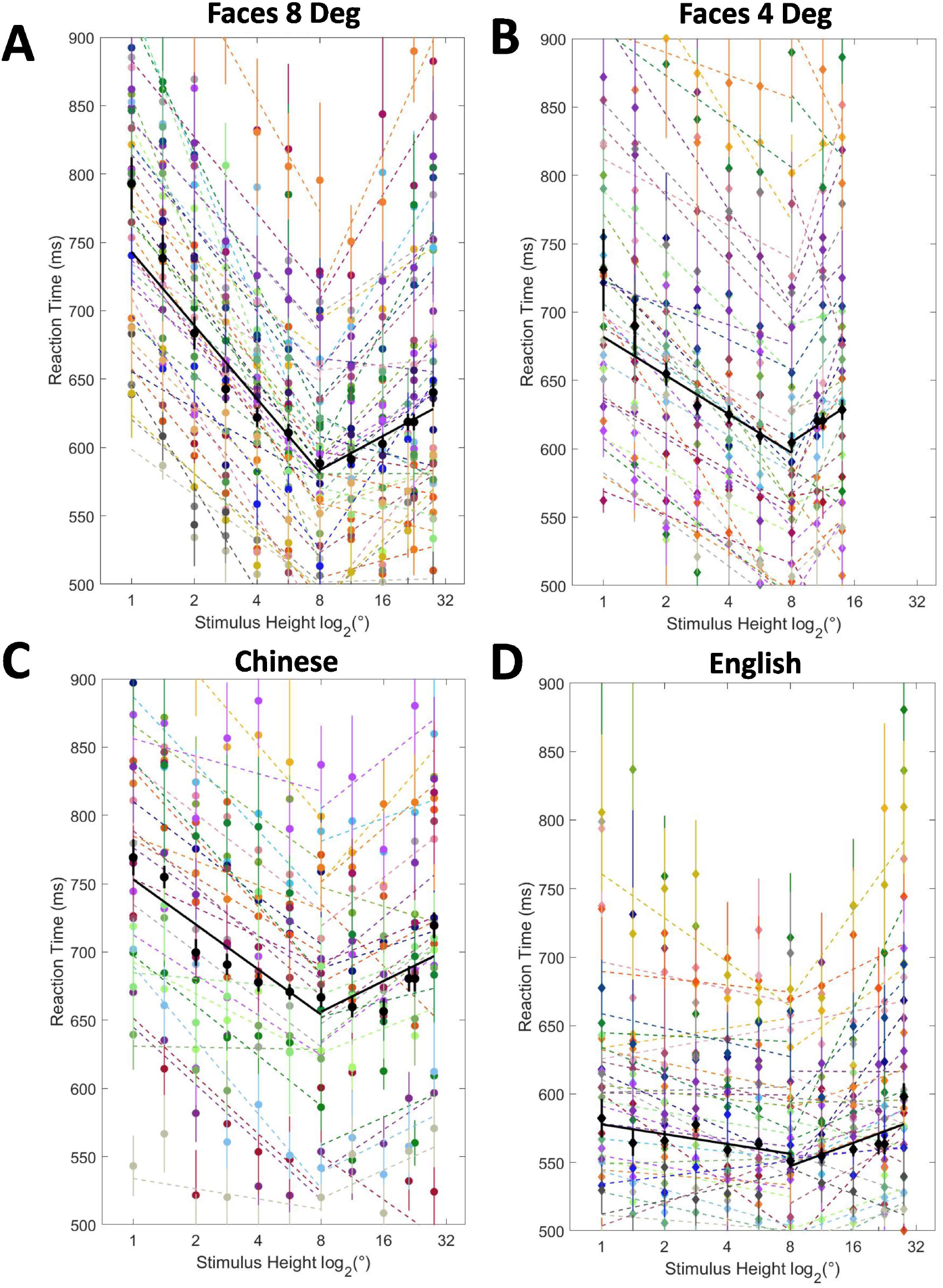
Raw data for every participant from Groups A, B, and D combined. These plots show raw data without any alterations. (A) 8 Degree Faces data, with 36 participants total. (B) 4 Degree Faces data, with 30 participants total. (C) 8 Degree Chinese data, 19 participants total. (D) 8 Degree English data, 23 participants total.

**Figure 2.**
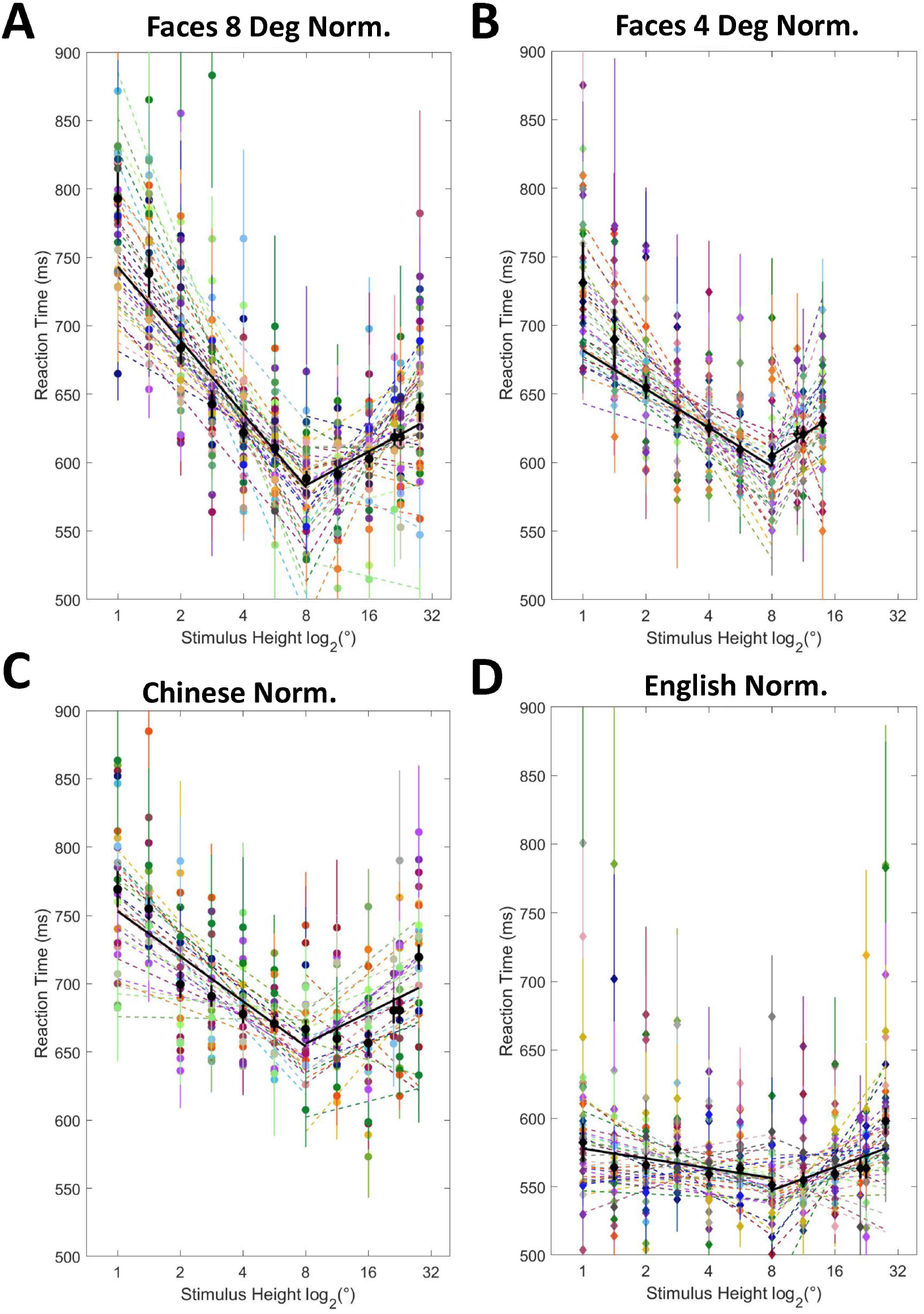
Normalized data for every participant from Groups A, B, and D combined. Each participant’s data was adjusted using the Cousineau method, shifting each point towards the aggregate line in black without affecting the slope. (A) 8 Degree Faces data, with 36 participants total. (B) 4 Degree Faces data, with 30 participants total. (C) 8 Degree Chinese data, 19 participants total. (D) 8 Degree English data, 23 participants total.

**Figure 3.**
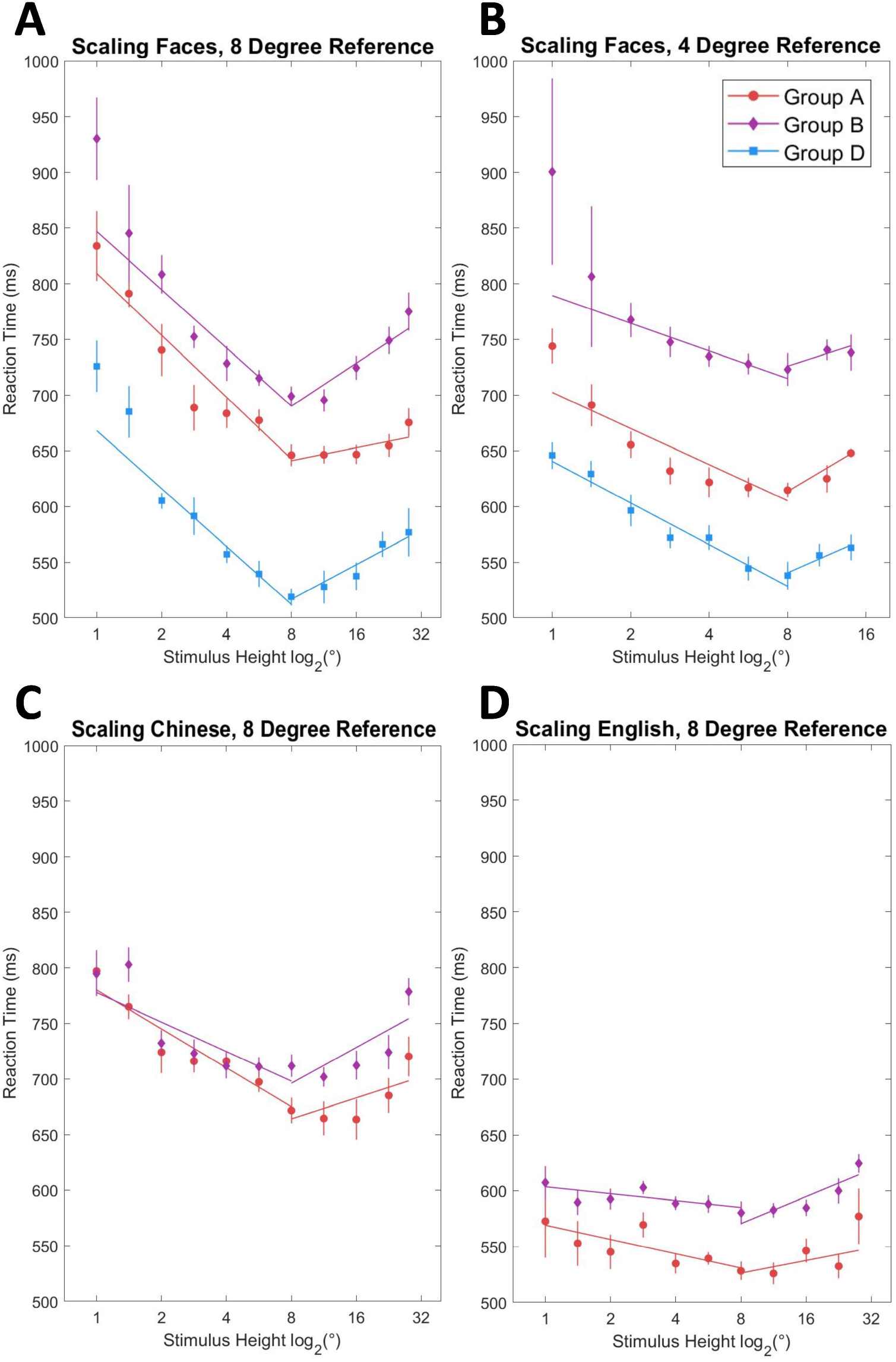
Comparison of 3 groups’ performances on the 4 experimental protocols. Groups A and D consisted of lab members not directly involved in this research topic. Group B was composed of UCLA undergraduate students that were recruited externally. Group B participants were tested using the Protocol A set, without any gaming incentives. Group D participants were given the Protocol B set, which did include a game aspect, as described in the Methods.

### 2.2 Separate Group Analysis

However, to verify the consistency of the data between the three independent data-taking groups, participants from each group needed to be separately evaluated, as in Figures 4 and 5. Figure 4 applies the same log scaled x-axis for this assessment, whereas Figure 5 utilizes the same data and converts the log degree height to the equivalent distance of an image from the position of the partcipant, using the formula 1.4 m x [8° / (Face height)]. For example, a stimulus height of 8 degrees corresponds to a distance of 1.4 m.

**Figure 4.**
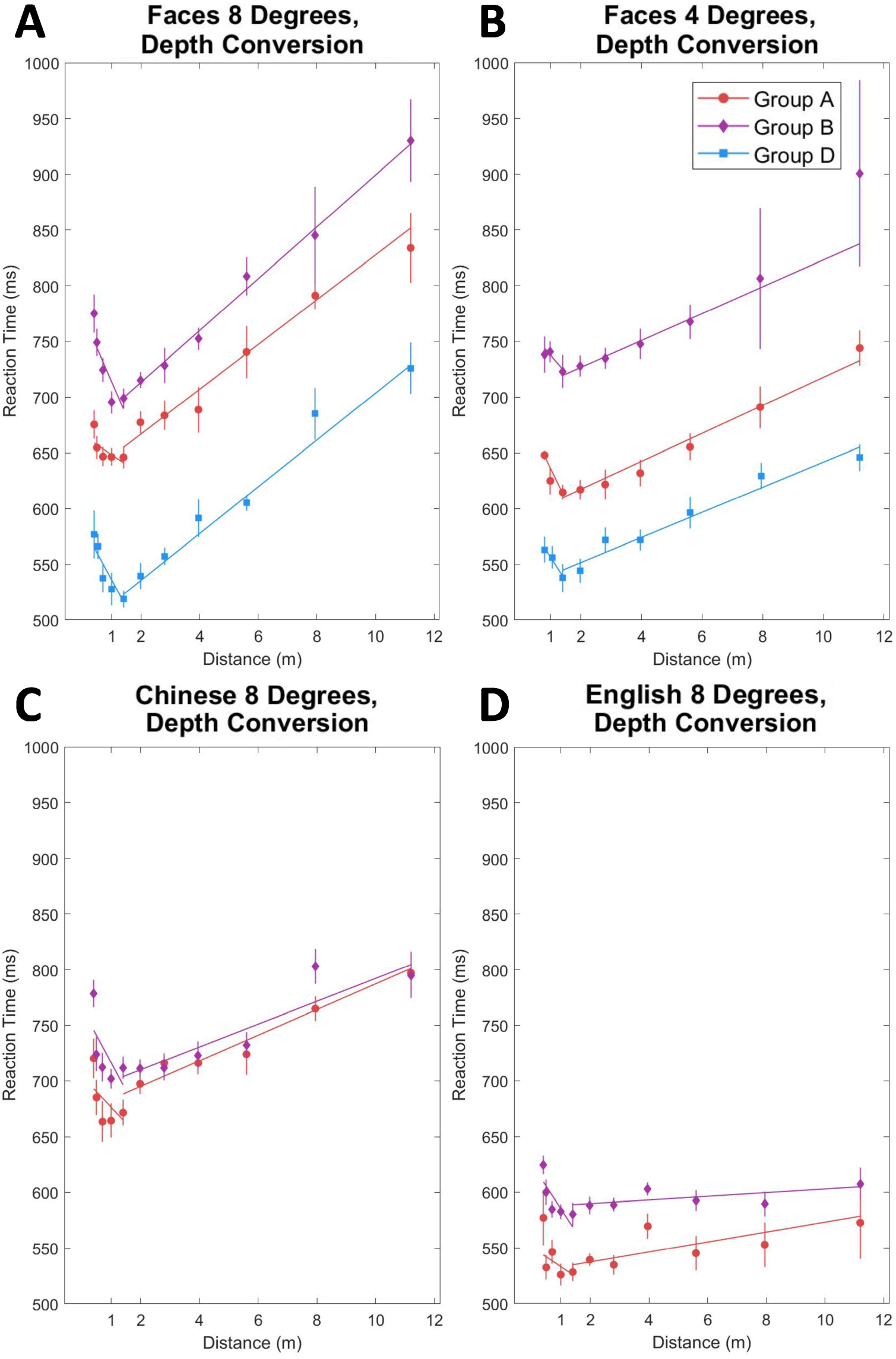
Comparison of 3 groups’ performances on 3 of the 4 protocols from Figure 3, now plotted with a depth conversion from stimulus height in degrees to distance in meters. (A, B) The x-axis for 8- and 4-degree face data has been converted to meters using 1.4 m x [8° / (Face height)], assuming a vertex at 1.4 m. (C, D) Character data was only tested with an 8-degree reference size.

**Figure 5.**
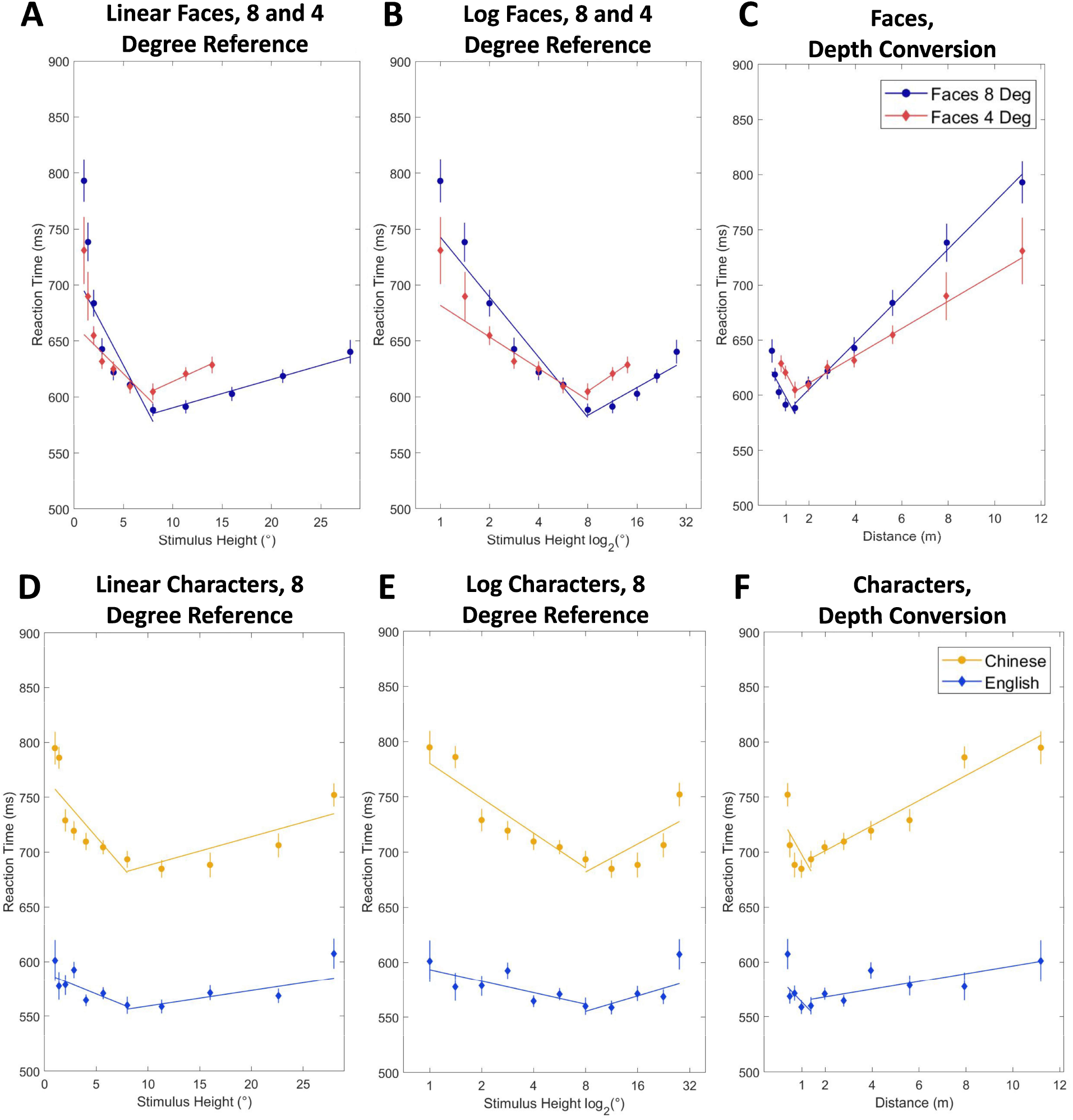
Aggregate analysis for all group data combined, including comparing faces memorized at 4- and 8-degree sizes. The fits were created assuming a vertex at 8 degrees, representing the hypothesis of a universal size for memorizing facial stimuli. (A, D) Data assessed with a linear scale for the x-axis. (B, E) X-axis was rescaled using log_2_(°). (C, F) For the depth conversion, the x-axis was transformed from stimulus height in degrees to the equivalent distance in meters. Small face sizes correlate to larger distances and vice versa.

In both figures, the slopes between the three groups are seen to be reasonably consistent, although the intercepts differ widely. For example, in the case of the 4-degree faces, Group B has an intercept that is about 150 ms slower than Group A. A likely explanation of this result is that Group B had little incentivization or motivation during data-taking, whereas Group A was composed of lab members and Group D was given a gamified protocol with high scores given for faster reaction times. Complete tables for the fit parameters of each group can be found under Appendix 1. Still, for the purposes of an analysis of the slopes and the appropriateness of a linear fit, the three groups’ data are suitable for aggregation.

### 2.3 Aggregate Analysis

The results of aggregating all participant data are displayed in Figure 5. Immediately evident in the log scale and depth conversion plots is that a linear fit can be reasonably applied to the data as a function of either stimulus height (in degrees) or distance (meters). However, a slight curve can still be observed in the log scale fit for small sizes (Figure 5B). The reduced chi-squared values evaluating the goodness of fit for these assumptions was calculated and is shown in Table 1. For small sizes, the distance conversion provides the smallest reduced chi squared values and therefore best encompasses the data. However, for large sizes, the linear scaled fit appears most appropriate. In both cases, the log scale provides an intermediate and acceptable reduced chi square.

**Table 1.**
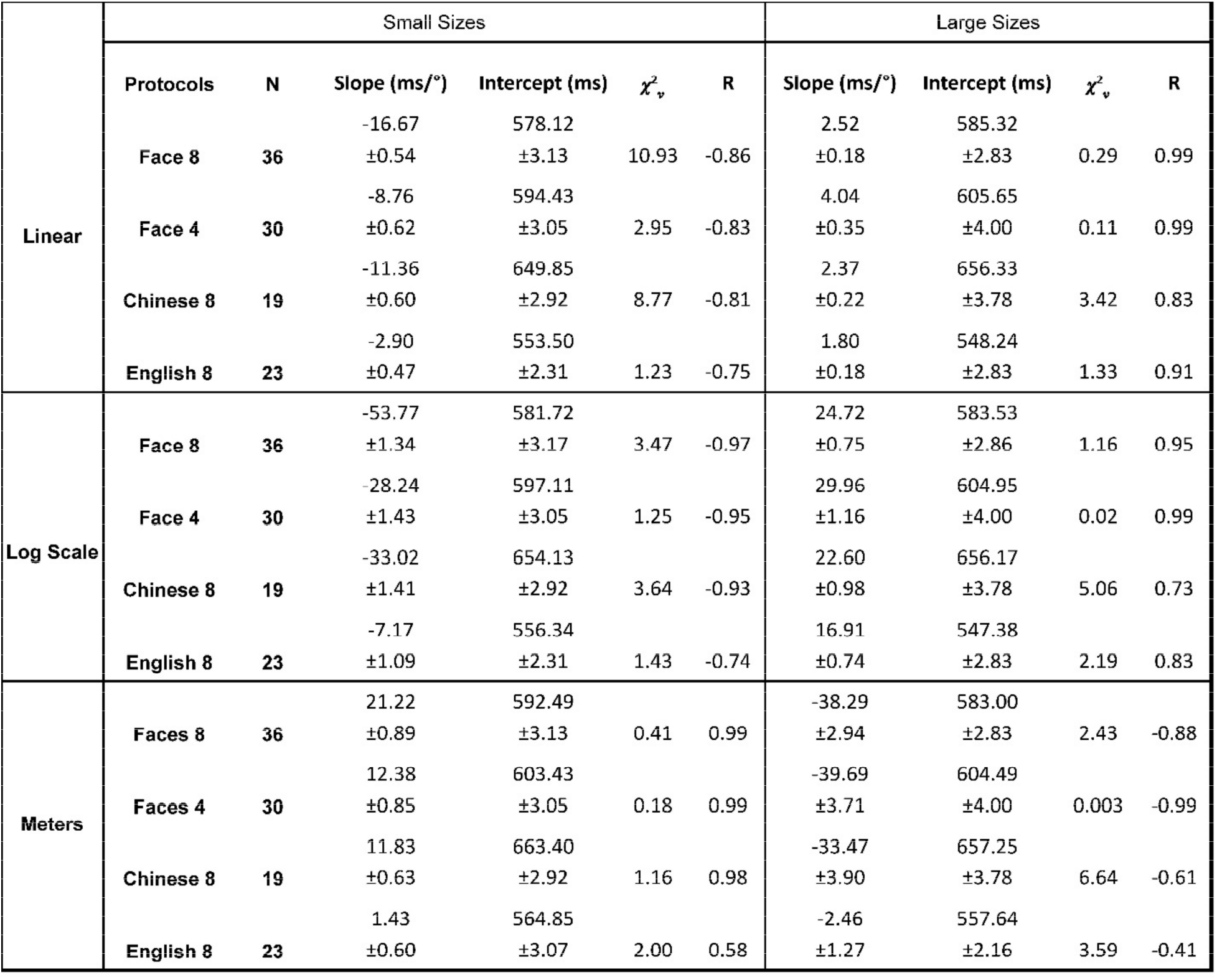
Fit parameters across all protocols, including slope and intercept for the regression lines, and reduced *χ*^2^(*χ*^2^*v*)and correlation coefficient R. Small size analyses refer to sizes smaller than or equal to the memorized size; large size fits were calculated with stimuli larger than or equal to the memorized size. The units for linear slope are in ms/°, log-scale is log_2_(ms/°), and meters is in ms/m.

In addition, the face data of Figure 5 shows that the fastest reaction time for each protocol is independent from the size memorized. Best visualized within Figure 5, the fastest response time was consistently observed at 8 degrees, even for experiments where the subject initially memorized a 4-degree face.

A comparison of the slopes and intercepts between the all 4 protocols are shown in Figure 6. Unfamiliar faces and Chinese data showed slower reaction times as intercepts at 600 ms or more were observed. Simpler and familiar characters, such as English, led to faster reaction times of about 550 ms. A similar trend was seen for the slopes of small size stimuli, but all protocols featured similar slopes for large sizes (sizes larger than the designated reference height).

**Figure 6.**
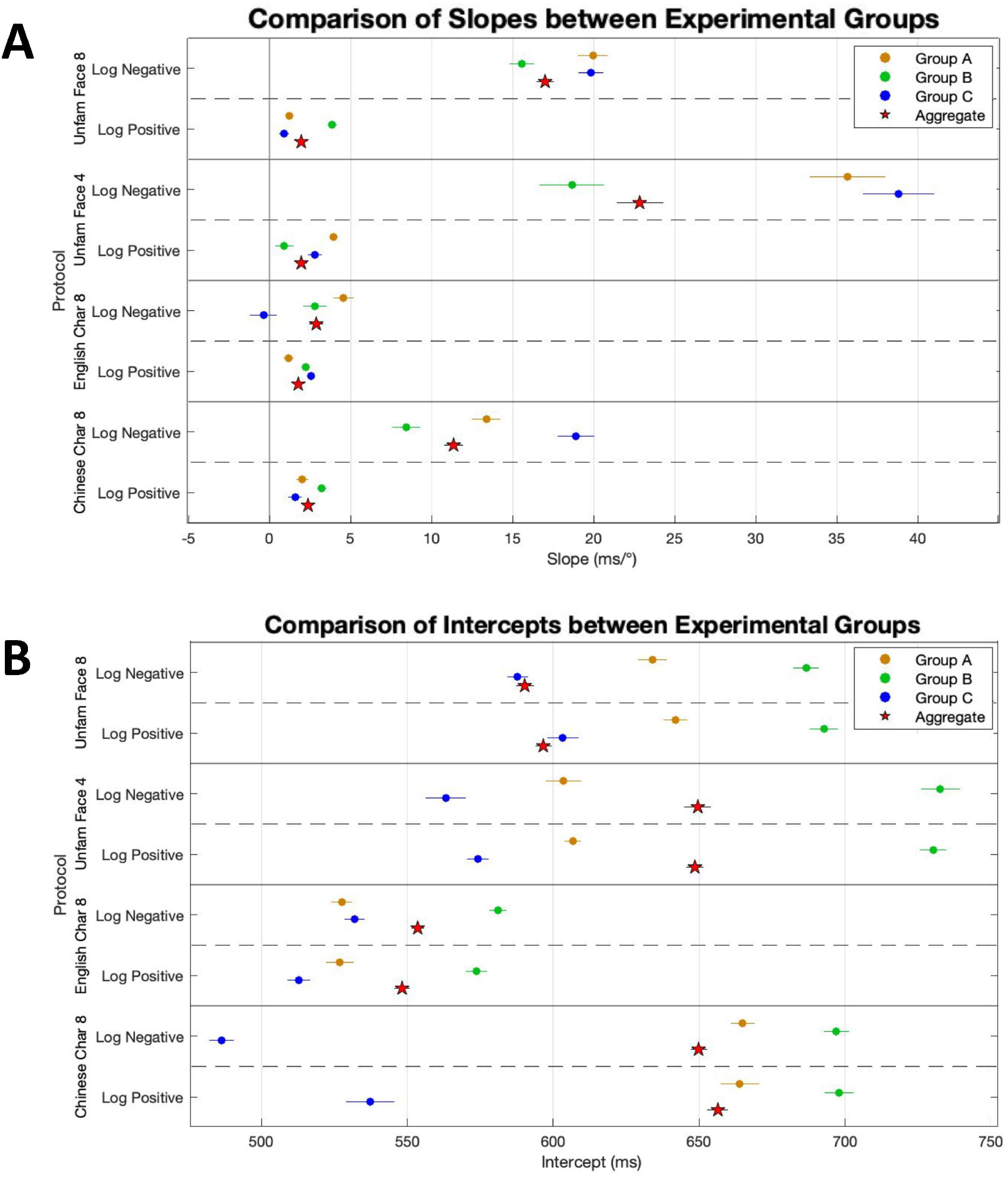
Slope and intercept plots for each of the four protocols when assessed with log-scaled stimulus height. (A) shows a comparison of slopes between each of the individual groups and the aggregate value. (B) compares intercepts between these groups and the aggregate.

## 3 Discussion

The results of these behavioral protocols encompass a wide range of visual stimuli, elucidating the mechanisms behind the encoding of an image and the transformations involved with a scaling factor. The objective of these protocols was to train a participant on a specific visual stimulus condition (i.e., size) and to test how a participant’s recognition reaction time is affected by changes in visual presentation from their original memorized condition. The encoding of a visual memory, in relation to its mapping onto the ventral pathway, ultimately relies on the concept of Hebbian plasticity (Hebb, 1949).

It is suspected that the scaling of visual information occurs at the V1 in a log-polar coordinate system, as the mapping of information onto the V1 cortex occurs in a log-polar manner (Abdollahi et al., 2014; Benson et al., 2014). The log-polar nature of the visual cortex is supported by retinotopic maps showing log-polar patterns of one-to-one mapping from the retina to the cortices (Arcaro et al.; Engel et al.). The results display strong evidence that image scaling, mapped onto the V1 Log-Polar Coordinate system, and top-down processing due to memory comparison heavily affects facial recognition. The fastest reaction times for facial recognition tended to correspond to sizes at or right above the reference size, portraying top-down processing. The participant compares the incoming stimulus with their memorized depiction, so it takes longer to respond to cues that appear more visually altered. In this case, cues that are increasingly larger or smaller than the memorized reference will take more time to react to.

The degree of difference from memory generates a symmetric decay in reaction time for sizes above and below the memorized reference. This is supported by the Left-Right linear regressions that meet at the reference size, their lowest RT value. The logarithmic mapping on the V1 is mimicked by the semi-log plot x-axis; the linear regressions for RT are appropriate when plotted with a logarithmic x-axis. These trends can be observed in the data collected, as the reduced chi squared values appear to be reasonably small for the left-fits (below reference size) on the semi log plots. One deviance from this is how the reduced chi squared values for the right-fits (above reference size) are superior for the linear scale plots. This could indicate that scaling up a memorized image displays less detrimental effects on reaction time, in comparison to scaling down.

Facial recognition has been thought to be processed by the right fusiform gyrus, as patients displaying lesions in this area have shown to have difficulty in recognizing facial features—or prosopagnosia (Barton et al., 2002). The current consensus of facial recognition calls for a more comprehensive model that incorporates a dominant top-down processing while describing the brain’s evolutionary features to understand scaling. In fact, a new model of vision proposes an extension to this current view, proposing that faces—especially unfamiliar ones—are processed by the ventral pathway in a top-down manner, in the V1 region. This localized processing at the V1 is specifically related to identification of objects at different sizes. Which leads to the relationship between facial recognition and scaling. In order to identify objects we have encountered, we must compare them with our memory of said object. A top-down mechanism indicates an object is recorded in our memory at a specific size. Therefore, the V1 becomes the center for incoming stimuli to be analyzed: where the memory of the object (at a particular size) is used to confirm the identity of what we see. In this study, this was demonstrated by reaction times, which showed that more processing time is required to identify an object that looks smaller or larger than the encoded memory.

As a result, the recognition of a face can be connected to its perceived distance from the observer, who must internally re-scale the stimulus to match the memorized reference size. If the stimulus height data is converted to equivalent distances and plotted as shown in in Figure 5C, a new linear regression can be fit. As Table 1 reports, these reduced chi-squared values are the smallest in comparison to the linear and log-scale analyses of the stimulus heights. Therefore, not only does the brain achieve scaling invariance via a log-polar mapping on V1, but it can simultaneously detect the distance of an object based on its size relative to a memorized reference (Bustanoby et al., 2022). This multi-tasking is essential to quickly understand an individual’s surroundings and recognize potential threats.

The evolutionary pressure for the observed mechanisms could have been the reliance of early humans in their vision for survival. In other words, differently from other mammals and their acute sense of smell and hearing, humans ended up developing a faster and more accurate way of determining objects’ size in their struggle to survive. The results support the assumption that the fastest reaction times will correspond to the object size of the participant’s first encounter with it. In line with this, the participant’s recognition speed decays as the object is further altered visually from its original state. Interestingly, this occurs in a mathematically symmetrical decay on both sides of the alteration spectrum; in decreasing and increasing sizes. Notably, this symmetry in reaction time decay corresponds to a logarithmic relationship between the change in scaling properties. These findings implicate the scaling presence in the V1 as a potential evolutionary advantage for fast recognition based on top-down memory processes of image comparison.

## 4 Materials and Methods

### 4.1 Participants

Participants were drawn from the undergraduate and graduate UCLA student population in accordance with approved procedures from the Institutional Review Board (IRB # 19-001799). Due to the COVID-19 pandemic, data-taking began remotely with internal members from the summer of 2020 until the summer of 2021. After which, standardized experimental setups were developed in-lab and external participants were recruited for additional, professional data-taking. These groups’ data were assessed separately for consistency, then combined in an aggregate analysis as shown in the Results.

The breakdown for the number of participants (n) for unfamiliar faces 8-degree reference is displayed in the table below. The faces 8-degree protocol had 36 data sets, faces 4-degree had 30, the letter protocol for familiar alphabets had 23, and Chinese had 19. The participants counts for each group under the remaining protocols can be found under Appendix 1.

There were three participant groups in total: the first was composed of internal lab members, while the others consisted of externally recruited participants. Both the group of internal lab members and the first group of external participants were tested on a non-gamified version of the protocol (Group A and B, respectively). The second group of external participants, Group D, was tested using the gamified version of the experimental protocol. As indicated in Table 1, data from each participant group was analyzed separately and then aggregated with the other groups for a final analysis containing all participants groups.

**Table 2.** Descriptions of the three independent data-taking groups, including participant count and whether a gamified protocol was used. *The participant count listed shows the distribution for 8 degree faces data.

| Group Letter | Source of Participants | Max n* | Equipment | Experimental Design |
| --- | --- | --- | --- | --- |
| A | Internal members from lab | 15 | 55" TV, with headrest | 3-choice Reaction Test, No game |
| B | External participants from Physics labs | 13 | 55" TV, with headrest | 3-choice Reaction Test, No game |
| D | External participants from Physics labs | 14 | 55" TV, with headrest | 3-choice Reaction Test, Game |
| Aggregate | Internal and external participants | 42 | 55" TV, with headrest | 3-choice Reaction Test |

Two versions, referred to as Protocol A and Protocol B, of experimental protocol were used in external data collection. To maximize the number of participants included in final analysis, data sets from both protocols were assessed. Protocol B was identical to Protocol A, with the addition of a video game-inspired “point system” element aimed towards motivating participants to achieve maximum facial recognition speed and accuracy. Both Protocols A and B relied on the same structure: a stimulus was displayed on screen, and the participant pressed an associated key in response. The visual stimuli studied included (1) unfamiliar faces, (2) English characters, and (3) Chinese characters. Protocols A and B were written in Python and performed using PsychoPy software. More details regarding the experimental procedures can be found under the Overview of Protocols.

### 4.2 Experimental Setup

Data for both protocols was taken with participants seated 1.4 meters (reference 8 degrees) or 2.8 meters (reference 4 degrees) away from a large television screen, with their eyes aligned with the center of the screen (Figure 7). Only Group D took data for 4- and 8-degree protocols while being seated at 1.4 meters for both. Participants’ head positions were kept stable using a custom-built, adjustable-height chin-rest. Distractions were minimized by way of cloth drapes placed on either side of the television screen.

**Figure 7.**
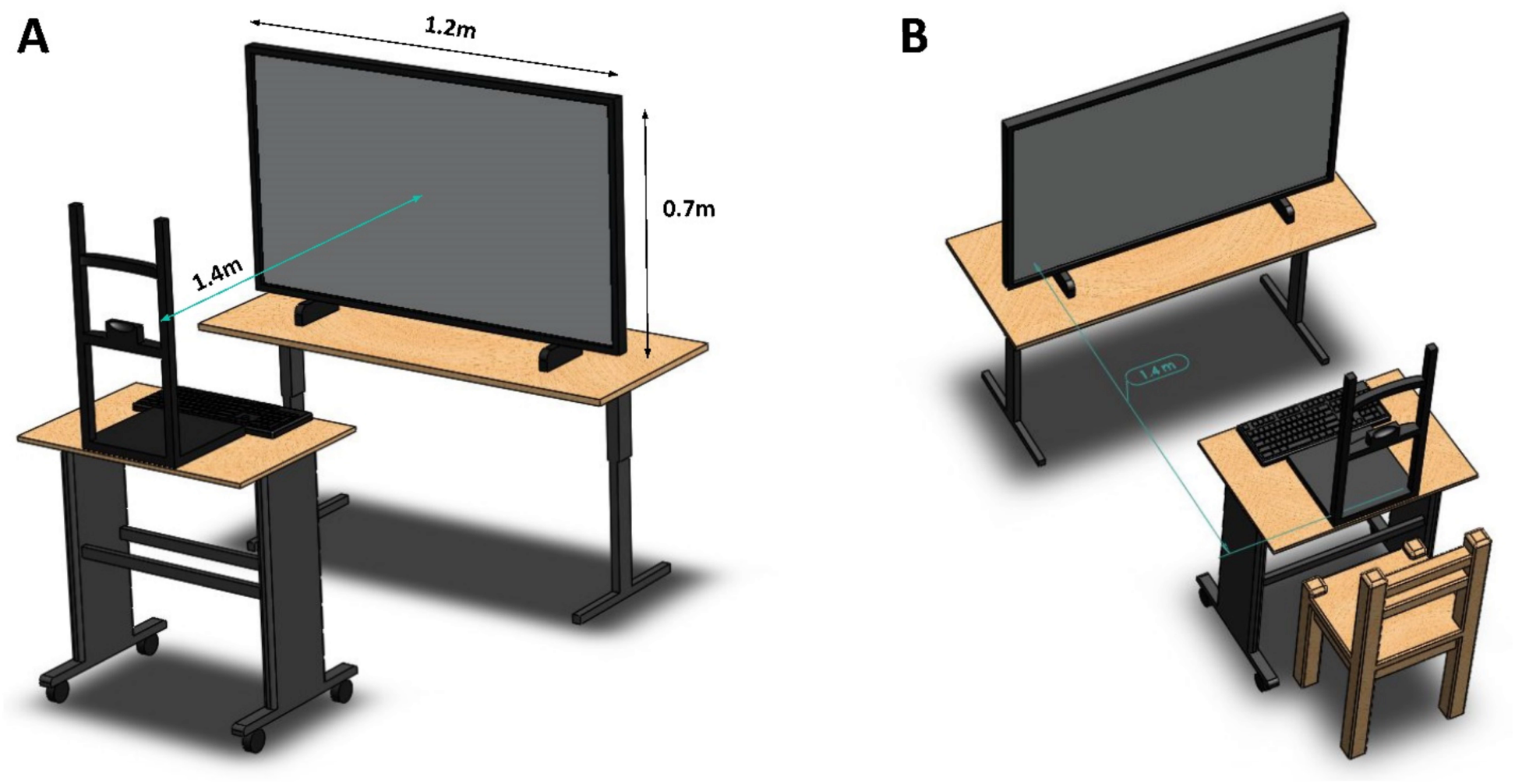
All participants sat 1.4 meters away from a 55 inch, 4k monitor for the 8 Degree Face, Chinese Character, and English Character experiments. Participants from Groups A and B sat 2.8 meters away from the screen for the 4 Degree Face experiment. Volunteers were instructed to rest their head on the chin rest and place three fingers of their dominant hand on the keys “v,”“b,” and “n.” Curtains around the system (not shown) provided additional privacy and limited distractions. (A,B) show alternate views of the system.

### 4.3 Equipment Calibration

The stimuli for the face scaling protocol were selected from online face databases and displayed at the center of the screen. All images had the same square dimensions with resolutions of 330×330 pixels. The protocols were coded in Python and performed using the PsychoPy platform. Each display screen was first calibrated through a calibration script, which measured the physical dimensions of the screen, distance of the participant to the screen, pixel densities, and more. These measurements were used in the experiment protocol in order to maintain regular dimensions of each stimulus across different systems. System input lag was also measured and calibrated for using a 960 Hz slow motion camera to record the time between an input signal and output image.

### 4.4 Protocols Overview

Protocol A began by displaying on-screen instructions indicating that participants would have thirty seconds to become familiarized with each visual stimuli displayed. Instructions were followed by a practice round in which visual stimuli were displayed at controlled reference sizes. These reference sizes were used for the practice to give participants the opportunity to encounter and learn visual stimuli at the specific initial conditions. In this practice round, the participant learned to associate different letters of the keyboard (e.g., “v,” “b,” or “n”) with each unique visual stimulus. All protocols were choice reaction time tests between 3 similar stimuli (3 CRT). For each trial, a stimulus appeared within a randomized interval of 600 to 1600 ms. This pause was selected to prevent stimuli from appearing within the refractory period of action potentials, while simultaneously preventing a fixed interstimulus time allowing participants to anticipate the next stimulus. Feedback was given after every key response: correct responses were indicated by a green outline around the image, and incorrect responses were indicated by red. The colored feedback was displayed for 1 second before the next trial began.

Following the initial practice round, Protocol A initiated an official data-taking round. Whereas practice rounds kept visual stimuli at predetermined constant sizes, data-taking rounds changed the scale of visual stimuli. However, Protocol A alternated between practice rounds and data taking rounds. There were a total of nine unique visual stimuli in the protocol organized in three sets, each set containing three similar images to be tested at a time. Throughout each protocol, the stimulus set changed twice to prevent participants from becoming overly familiar with the tested stimuli. Experiments continued for 150 trials with 15 trials per stimulus height, on average. For 8-degree reference protocols, 11 unique stimulus heights were tested ranging from 1 to 28 degrees. For 4-degree protocols, 9 unique heights were tested from 1 to 14 degrees. Mistaken responses were recorded and retested at the end of the experiment.

Protocol B was identical to Protocol A, with the addition of a video game-inspired “point system” element aimed towards motivating participants to achieve maximum facial recognition speed and accuracy. Correct responses netted participants an arbitrary number of points, and after completing the protocol, participants were able to see their final scores in relation to the scores achieved by past participants.

Both protocols could be run using (1) Unfamiliar faces, (2) English characters, or (3) Chinese characters as visual stimuli. Unfamiliar faces were studied using reference sizes of both 4 and 8 degrees (with the participant distanced 2.8 m and 1.4 m from the screen, respectively), while English and Chinese characters were tested using only a reference size of 8 degrees (with the participant distanced 1.4 m from the screen).

**Figure 7.**
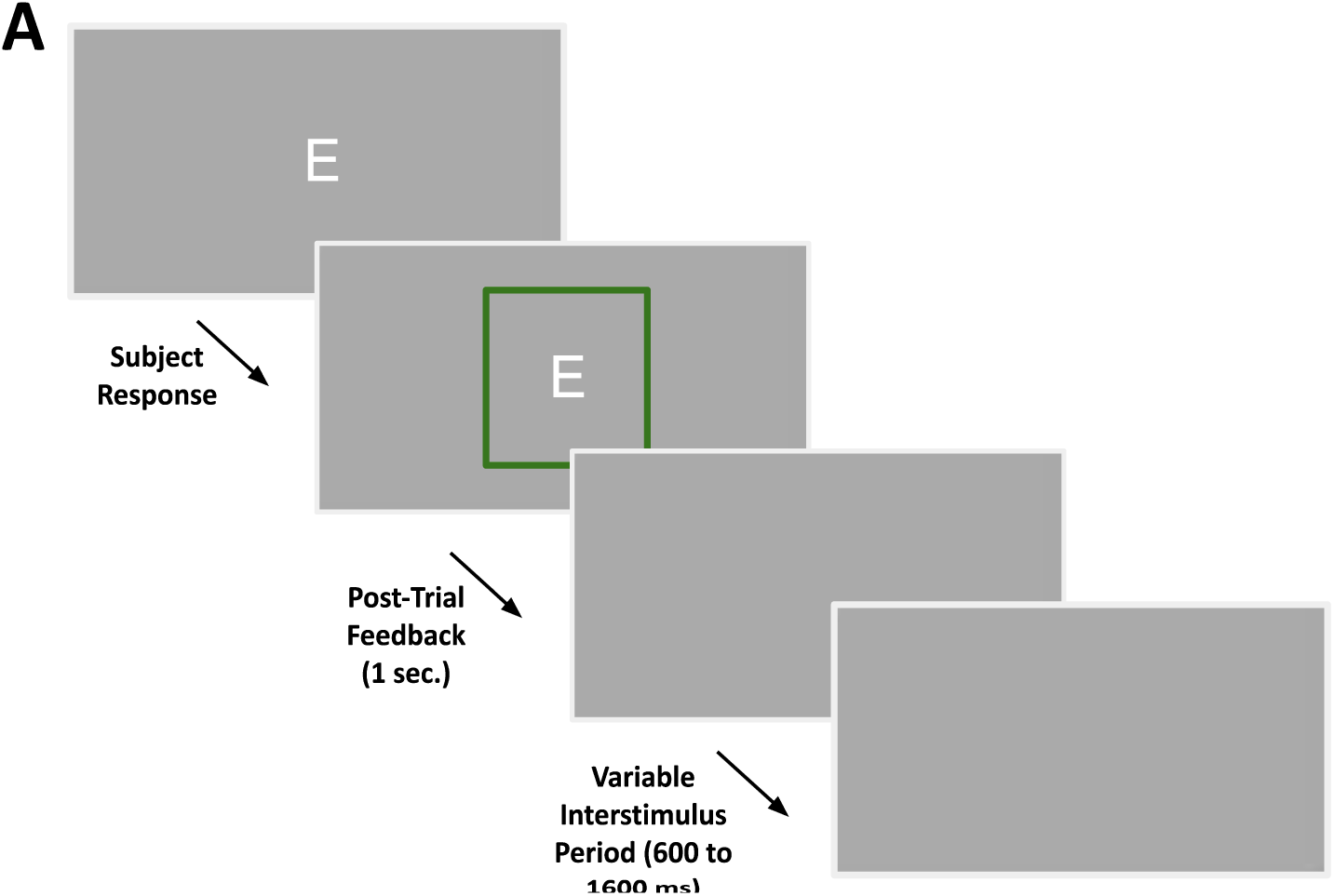
Outline of the experimental procedures for a general scaling experiment. (A) Character stimulus example. Following a participant response, a green or red outline appeared around the stimulus to indicate a correct or incorrect input, respectively. The outline was kept for 1 second before disappearing. A variable interstimulus period of 600 to 1600 ms followed, until the next stimulus appeared to begin the next trial.

**Figure 8.**
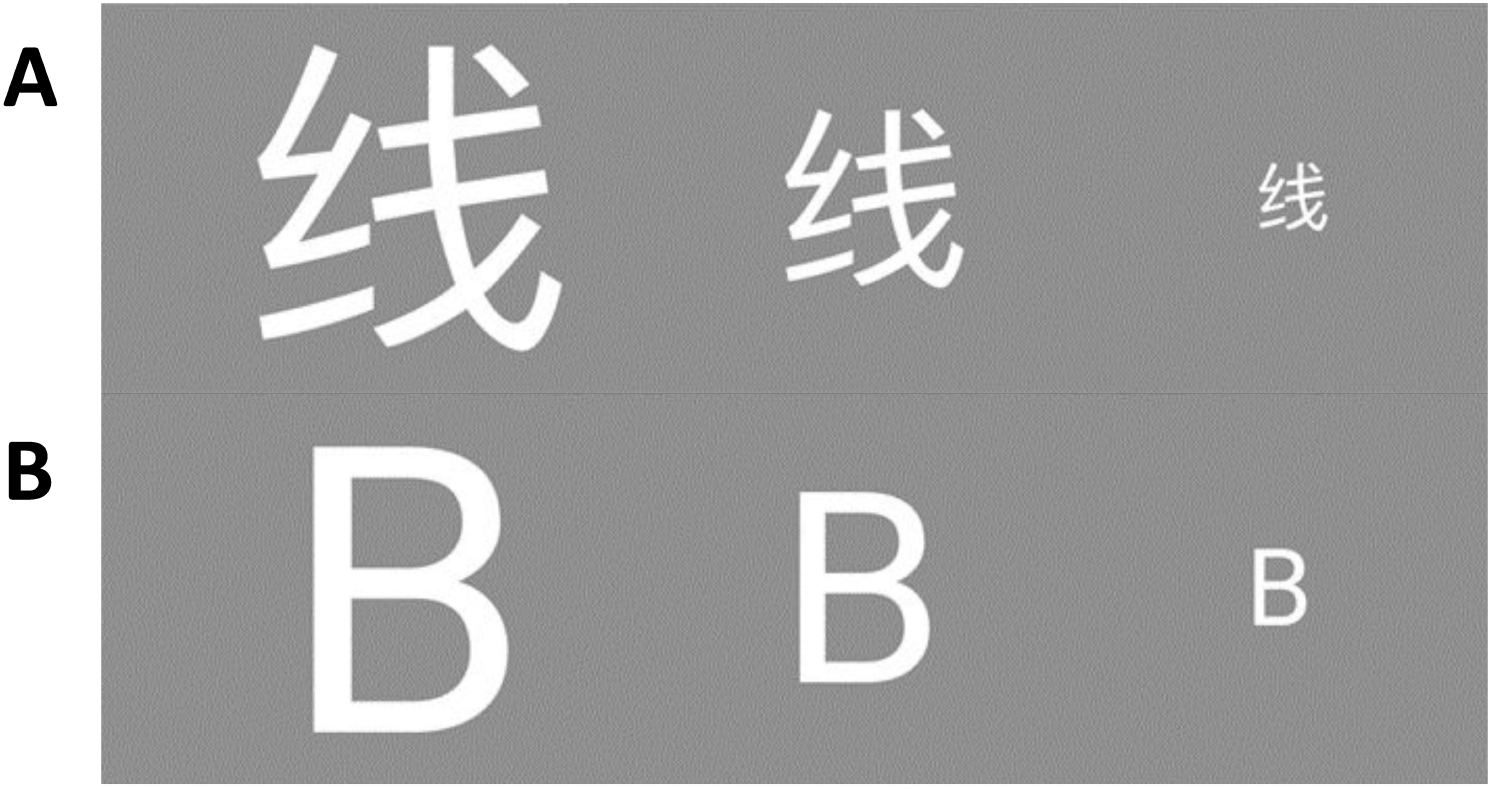
Scaling examples for each of the three types of tested stimuli. Each experiment investigated how the reaction time was affected in response to stimuli presented at varying sizes. Participants were presented with (A) Chinese characters, (B) English characters, and unfamiliar faces. The height of the stimuli ranged from 1 to 28 degrees.

### 4.5 Statistical Analysis

Reaction time trial data were categorized by their stimulus heights, and further analysis was done within the data separated by each height.

#### Outlier Analysis

Outlier data were removed before averaging the RT at each eccentricity. Data points that fell below the 25th percentile minus 1.5 times the interquartile range or above the 75th percentile plus 1.5 times the interquartile range for reaction time data at each angle were removed. The resulting data points were then averaged and used in chi square calculations.

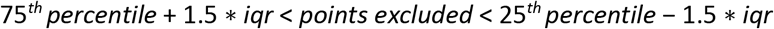

#### Error Analysis

The resulting reaction times were averaged, and the error bars were plotted using the calculated standard error, taking the standard deviation of the eccentricity angle reaction time divided by the square root of the number of trials within the eccentricity angles. After reaction time data was categorized by angle and outliers were removed, the standard deviation and standard error were calculated, with the standard error being used in the plots.

#### Hypothesis Testing

To ensure that an attention shift occurred during each choice RT block, response accuracy was analyzed using a one sample t-test for a mean of 1, with 3 being the number of available choices for each protocol. The test was performed for each stimulus height and participant’s data was removed for a given height if the corresponding p-value was greater than 0.05.

#### Normalization of average RT

To better represent the relationship between RT and stimulus height in the aggregated analysis, we normalize each participant’s data to a global average for each protocol. We normalize the data by subtracting the appropriate participants’ mean performance from each observation, and then add the grand mean score to every observation. Let y be the i-th participants score in the j-th condition (i =1,… N and j = 1,… M). Then define the normalized observations z-

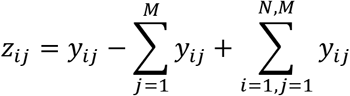

#### Line of best fit and correlation coefficient (χ^2^)

A best fit line was added using a least-squares linear regression for both positive and negative heights and their corresponding reaction times, with the memorized reference size being included in both positive and negative fits. The linear regression returned slope and intercept parameters as well as the correlation coefficient and a two-sided p-value in which the null hypothesis assumes a slope of 0. Additionally, the standard error of the fit was returned.

A best fit line of the form y = mx + b was calculated using chi square minimization to find the best fit slope and intercept parameters for both positive and negative heights with each side including the reference height. From the chi square minimization, error estimates for the slope and intercept parameters were computed by holding the other parameter constant at the best fit and calculating which parameter values would yield the minimum chi square value + 1. This process was done for both slope and intercept parameters. The reduced chi square was also calculated to determine goodness of fit by dividing the minimum chi square value by the degrees of freedom. The correlation coefficient was also calculated using the Numpy library in Python.

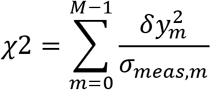

where 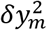 is defined as the mth residual 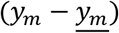 and *σ_meas,m_* is the standard error of the reaction time data for an angle m.

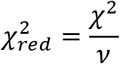

where *v* is the degrees of freedom.

## Notes about Authors and Data

### Data availability

The stimuli and code to calibrate the monitors, analyze data, and run the experiments can be found at https://github.com/brianta1/ScalingRT.

## Acknowledgements

A total of ~50 UCLA students helped us as participants in various stages of RT experiments in 2020 – 2022. We thank them for their kind commitment.

This work was in part supported by the Dean’s office of life science, Dean’s office of physical science, Chair’s office of the department of physics and astronomy, and the Instructional Improvement grant by the Center for the Advancement of Teaching, all at the University of California, Los Angeles.

## Author contributions

KA developed the concept and supervised the project. BT and PW designed and developed the experiments. DE managed the IRB regulations. All listed authors coordinated the data taking. BT, MS, and VL analyzed the data. BT, AS, AD, and MS wrote the original manuscript. KA and AB edited and proofread.

