## Supplemental Information for "Visual Perception of 3D Space and Shape in Time - Part III 2D Shape Recognition by Log-Scaling"

| Linear Scale | | | | | | | | | | |
|---|---|---|---|---|---|---|---|---|---|---|
| | | Small Sizes | | | | Large Sizes | | | | |
| | Group | Slope (ms/m) | Intercept (ms) | $\chi^2_\nu$ | R | Slope (ms/m) | Intercept (ms) | $\chi^2_\nu$ | R | N |
| **Face 8** | **A** | -19.97 | 634.08 | 4.13 | -0.86 | 1.21 | 641.79 | 0.41 | 0.88 | 15 |
| | **B** | -15.57 | 686.61 | 4.92 | -0.84 | 3.88 | 692.75 | 0.55 | 0.98 | 13 |
| | **C** | -19.84 | 587.80 | 5.73 | -0.87 | 0.91 | 603.28 | 0.46 | 0.72 | 7 |
| | **Aggregate** | -17.02 | 590.33 | 10.24 | -0.86 | 1.96 | 596.66 | 0.63 | 0.96 | 35 |
| **Face 4** | **A** | -35.67 | 603.51 | 3.43 | -0.89 | 3.95 | 606.67 | 1.26 | 0.78 | 6 |
| | **B** | -18.66 | 732.60 | 0.76 | -0.84 | 0.92 | 730.16 | 0.37 | 0.52 | 10 |
| | **C** | -38.83 | 563.11 | 0.38 | -0.97 | 2.81 | 574.22 | 2.14 | 0.88 | 5 |
| | **Aggregate** | -36.87 | 639.95 | 8.39 | -0.86 | 2.13 | 648.75 | 0.25 | 0.94 | 21 |
| **Chinese 8** | **A** | -13.39 | 664.92 | 2.02 | -0.88 | 2.02 | 663.93 | 0.84 | 0.84 | 8 |
| | **B** | -8.46 | 696.97 | 4.58 | -0.74 | 3.23 | 697.93 | 2.29 | 0.85 | 11 |
| | **C** | -18.90 | 486.31 | 3.84 | -0.75 | 1.59 | 537.29 | 1.36 | 0.50 | 4 |
| | **Aggregate** | -11.36 | 649.85 | 8.77 | -0.81 | 2.37 | 656.33 | 3.42 | 0.83 | 23 |
| **English 8** | **A** | -4.56 | 527.56 | 0.95 | -0.76 | 1.17 | 526.82 | 1.15 | 0.79 | 12 |
| | **B** | -2.80 | 580.99 | 0.64 | -0.74 | 2.25 | 573.67 | 0.70 | 0.94 | 15 |
| | **C** | 0.35 | 531.86 | 1.39 | -0.004 | 2.58 | 512.92 | 2.16 | 0.71 | 6 |
| | **Aggregate** | -2.90 | 535.50 | 1.23 | -0.75 | 1.80 | 548.24 | 1.33 | 0.91 | 33 |

**Table A1.** Individual group and aggregate parameters for linear scale data across all 4 protocols.

| Log Scale | | | | | | | | | | |
|---|---|---|---|---|---|---|---|---|---|---|
| | Small Sizes | | | | | Large Sizes | | | | |
| | **Group** | **Slope (ms/m)** | **Intercept (ms)** | $\chi^2_\nu$ | **R** | **Slope (ms/m)** | **Intercept (ms)** | $\chi^2_\nu$ | **R** | **N** |
| **Face 8** | **A** | -55.61 | 642.54 | 1.33 | -0.96 | 11.68 | 641.11 | 0.65 | 0.80 | 15 |
| | **B** | -52.26 | 690.10 | 2.08 | -0.96 | 38.61 | 690.13 | 1.31 | 0.95 | 13 |
| | **C** | -56.47 | 602.79 | 2.80 | -0.96 | 8.97 | 602.89 | 0.54 | 0.62 | 7 |
| | **Aggregate** | -54.10 | 594.30 | 3.39 | -0.96 | 19.23 | 595.36 | 1.46 | 0.9 | 35 |
| **Face 4** | **A** | -59.86 | 607.90 | 1.19 | -0.97 | 23.16 | 602.88 | 2.24 | 0.66 | 6 |
| | **B** | -39.43 | 732.46 | 0.48 | -0.92 | 3.97 | 730.37 | 0.42 | 0.41 | 10 |
| | **C** | -61.01 | 565.64 | 0.51 | -0.97 | 15.80 | 570.95 | 1.79 | 0.88 | 5 |
| | **Aggregate** | -70.57 | 640.21 | 3.11 | -0.97 | 10.23 | 648.16 | 0.48 | 0.86 | 21 |
| **Chinese 8** | **A** | -35.00 | 675.11 | 0.84 | -0.96 | 18.93 | 664.12 | 1.22 | 0.74 | 8 |
| | **B** | -26.56 | 698.12 | 2.75 | -0.87 | 32.22 | 696.18 | 3.66 | 0.76 | 11 |
| | **C** | -44.49 | 504.42 | 1.37 | -0.90 | 14.49 | 537.99 | 1.55 | 0.40 | 4 |
| | **Aggregate** | -33.02 | 654.13 | 3.64 | -0.93 | 22.60 | 656.17 | 5.06 | 0.73 | 23 |
| **English 8** | **A** | -12.80 | 530.69 | 0.98 | -0.77 | 11.32 | 526.13 | 1.20 | 0.73 | 12 |
| | **B** | -6.17 | 584.99 | 0.76 | -0.72 | 24.23 | 570.50 | 1.49 | 0.87 | 15 |
| | **C** | 1.17 | 531.98 | 1.38 | 0.04 | 27.68 | 511.40 | 3.45 | 0.63 | 6 |
| | **Aggregate** | -7.17 | 556.34 | 1.43 | -0.74 | 16.91 | 547.38 | 2.19 | 0.83 | 33 |

**Table A2.** Individual group and aggregate parameters for log scale data across all 4 protocols.

| Depth Conversion (Meters) | | | | | | | | | | |
|---|---|---|---|---|---|---|---|---|---|---|
| | Small Sizes | | | | | Large Sizes | | | | |
| | Group | Slope (ms/m) | Intercept (ms) | $\chi^2_\nu$ | R | Slope (ms/m) | Intercept (ms) | $\chi^2_\nu$ | R | N |
| **Face 8** | **A** | 20.08 | 654.92 | 0.65 | 0.99 | -17.64 | 641.17 | 0.88 | -0.70 | 15 |
| | **B** | 23.24 | 699.43 | 0.21 | 1.00 | -60.36 | 689.41 | 2.50 | -0.88 | 13 |
| | **C** | 19.25 | 626.77 | 3.24 | 0.99 | -14.25 | 602.73 | 0.61 | -0.53 | 7 |
| | **Aggregate** | 21.25 | 604.99 | 0.45 | 1.00 | -29.54 | 595.09 | 2.48 | -0.82 | 35 |
| **Face 4** | **A** | 14.80 | 616.84 | 0.09 | 1.00 | -21.11 | 600.49 | 3.63 | -0.54 | 6 |
| | **B** | 13.91 | 733.07 | 0.17 | 0.98 | -2.51 | 730.96 | 0.46 | -0.28 | 10 |
| | **C** | 14.63 | 569.98 | 1.48 | 0.92 | -14.36 | 568.02 | 1.47 | -0.87 | 5 |
| | **Aggregate** | 14.81 | 652.69 | 0.19 | 0.99 | -7.76 | 648.21 | 0.74 | -0.75 | 21 |
| **Chinese 8** | **A** | 11.52 | 688.22 | 0.90 | 0.97 | -27.84 | 665.13 | 1.57 | -0.63 | 8 |
| | **B** | 10.26 | 703.94 | 1.44 | 0.92 | -49.24 | 696.40 | 5.21 | -0.65 | 11 |
| | **C** | 14.85 | 519.67 | 0.58 | 0.97 | -19.95 | 540.08 | 1.72 | -0.30 | 4 |
| | **Aggregate** | 11.83 | 663.40 | 1.16 | 0.98 | -33.47 | 657.25 | 6.64 | -0.61 | 23 |
| **English 8** | **A** | 4.48 | 534.62 | 1.20 | 0.75 | -17.70 | 525.81 | 1.26 | -0.67 | 12 |
| | **B** | 1.63 | 588.77 | 0.92 | 0.69 | -40.45 | 568.66 | 2.54 | -0.78 | 15 |
| | **C** | -0.21 | 530.95 | 1.39 | -0.01 | -45.59 | 512.19 | 4.94 | -0.56 | 6 |
| | **Aggregate** | 1.43 | 564.85 | 2 | 0.58 | -2.46 | 557.64 | 3.59 | -0.41 | 33 |

**Table A3.** Individual group and aggregate parameters for depth conversion data across all 4 protocols.

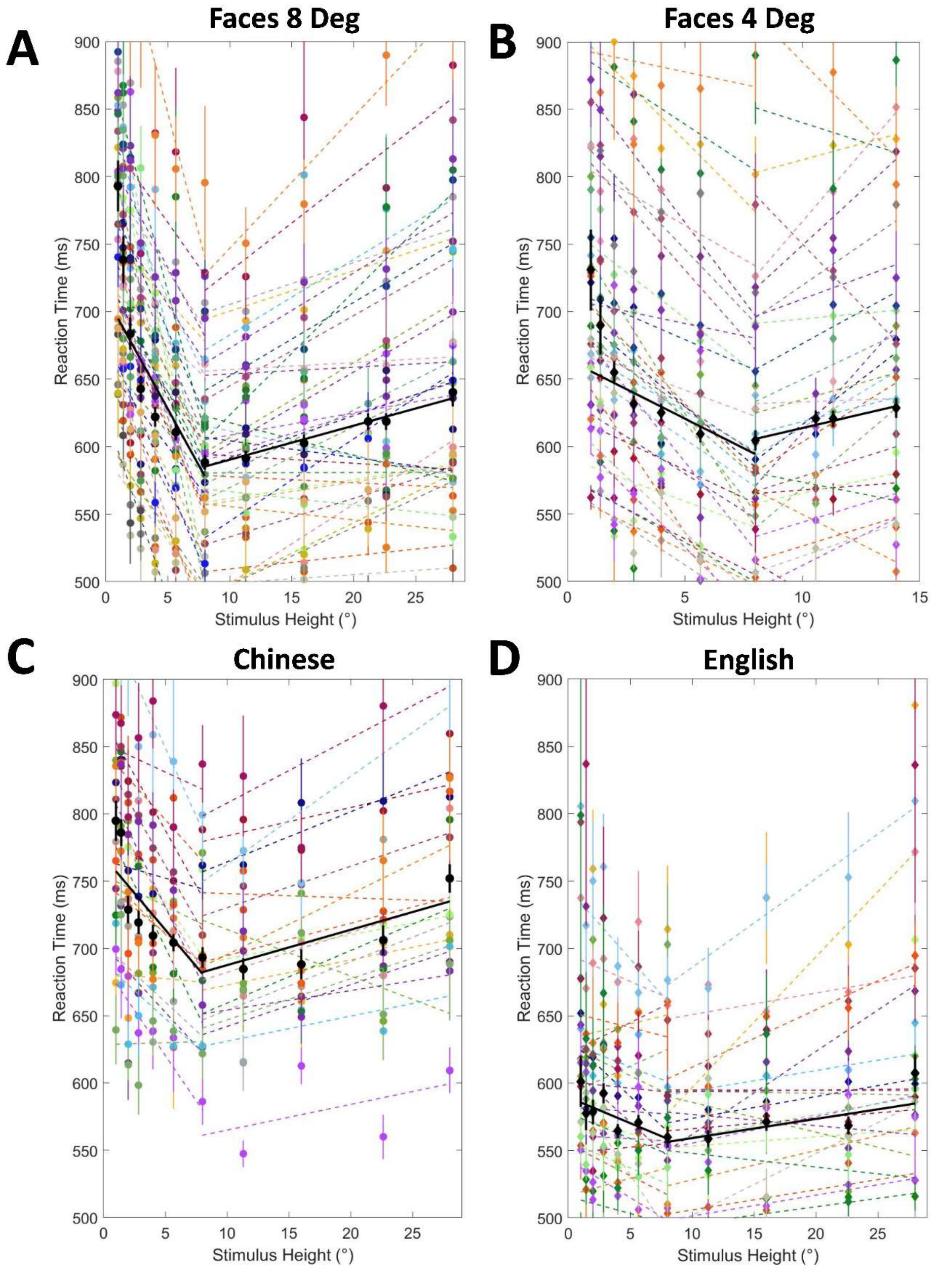

**Figure A1.** Linear-scale data for every participant from Groups A, B, and D combined. These plots show raw data without any alterations. (A) 8 Degree Faces data, with 36 participants total. (B) 4 Degree Faces data, with 30 participants total. (C) 8 Degree Chinese data, 19 participants total. (D) 8 Degree English data, 23 participants total.

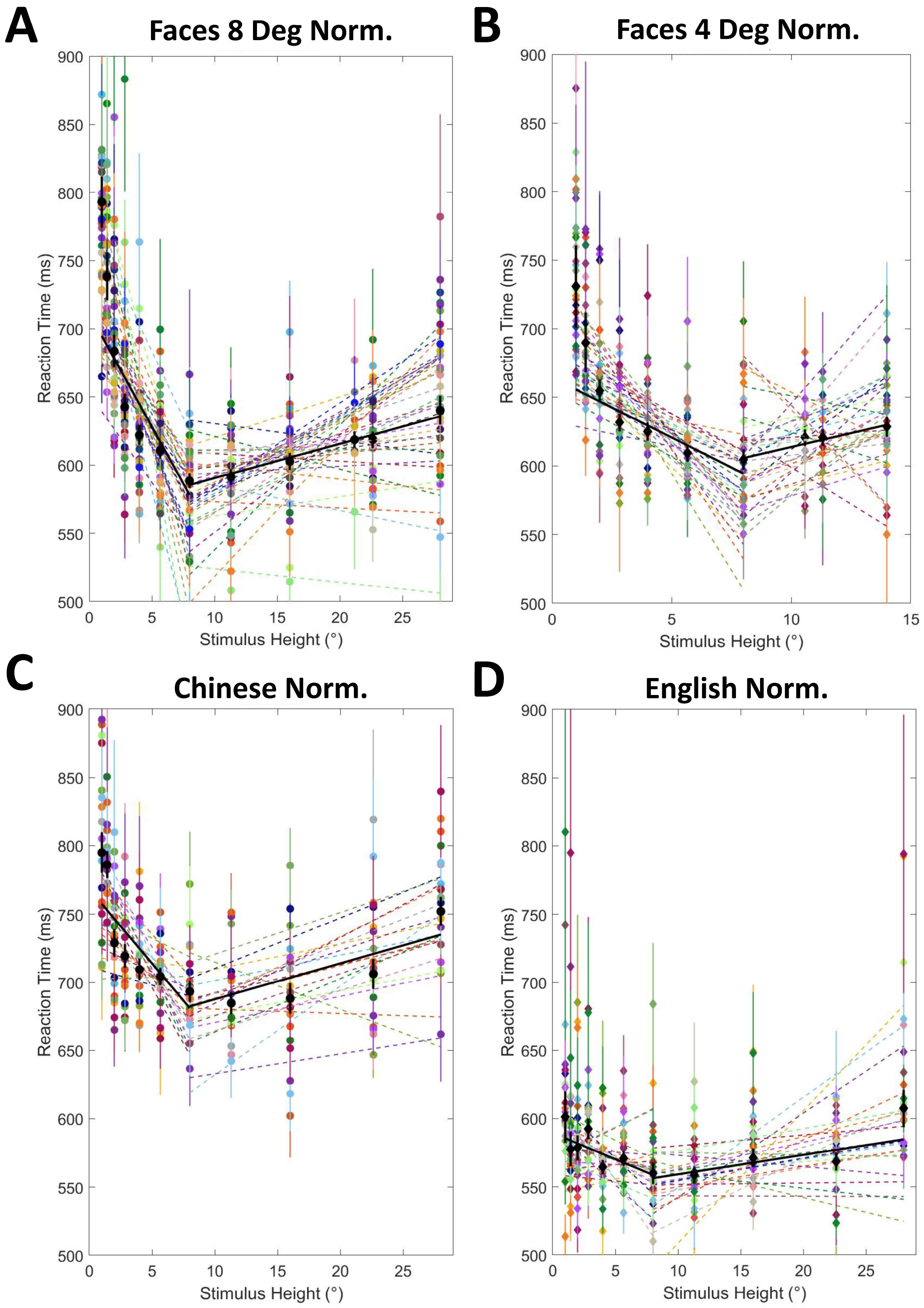

**Figure A2.** Linear-scale data for every participant from Groups A, B, and D combined. These plots show normalized data without affecting the slopes. (A) 8 Degree Faces data, with 36 participants total. (B) 4 Degree Faces data, with 30 participants total. (C) 8 Degree Chinese data, 19 participants total. (D) 8 Degree English data, 23 participants total.

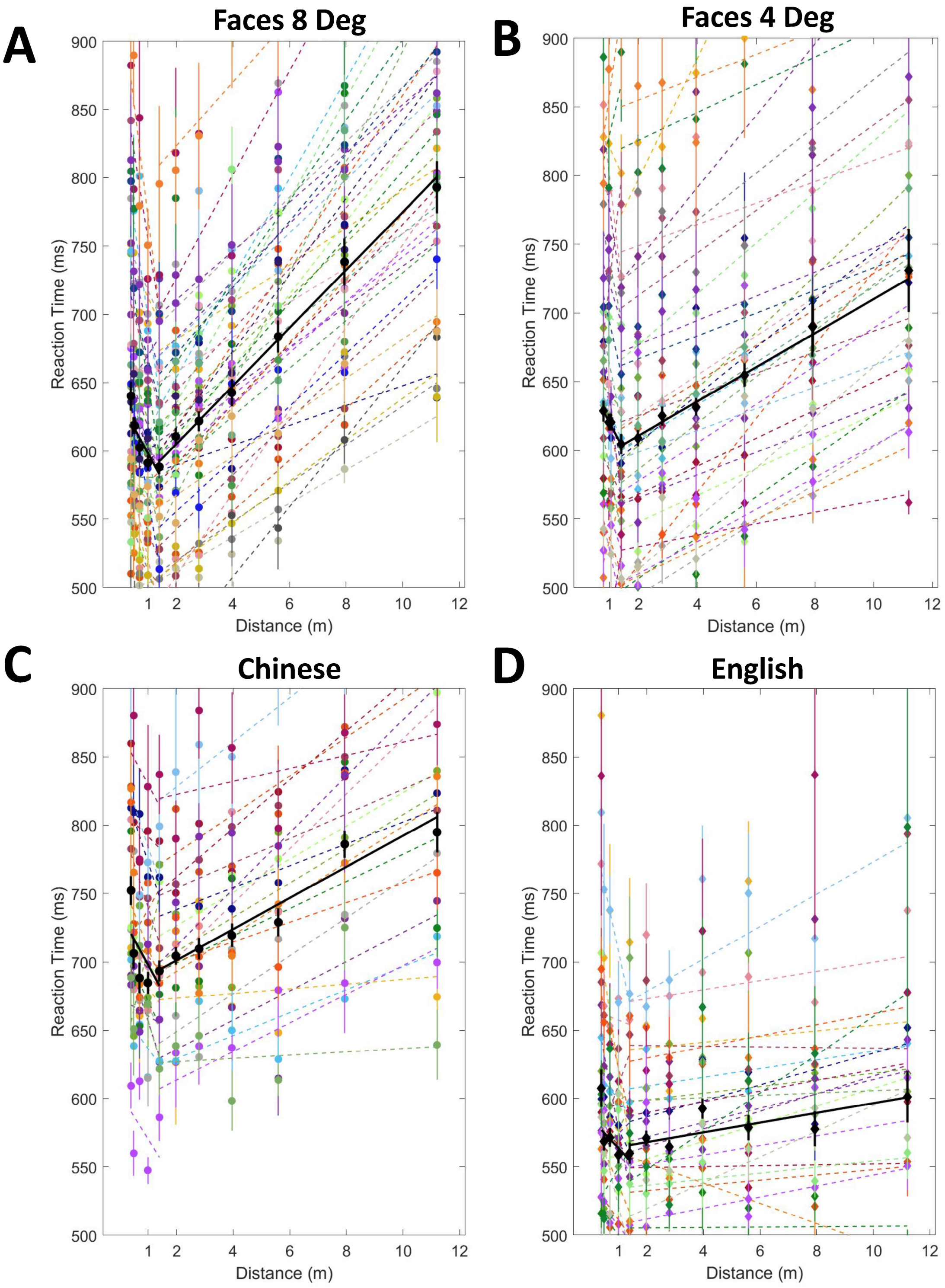

**Figure A3.** Distance conversion data for every participant from Groups A, B, and D combined. These plots show raw data without any alterations. (A) 8 Degree Faces data, with 36 participants total. (B) 4 Degree Faces data, with 30 participants total. (C) 8 Degree Chinese data, 19 participants total. (D) 8 Degree English data, 23 participants total.

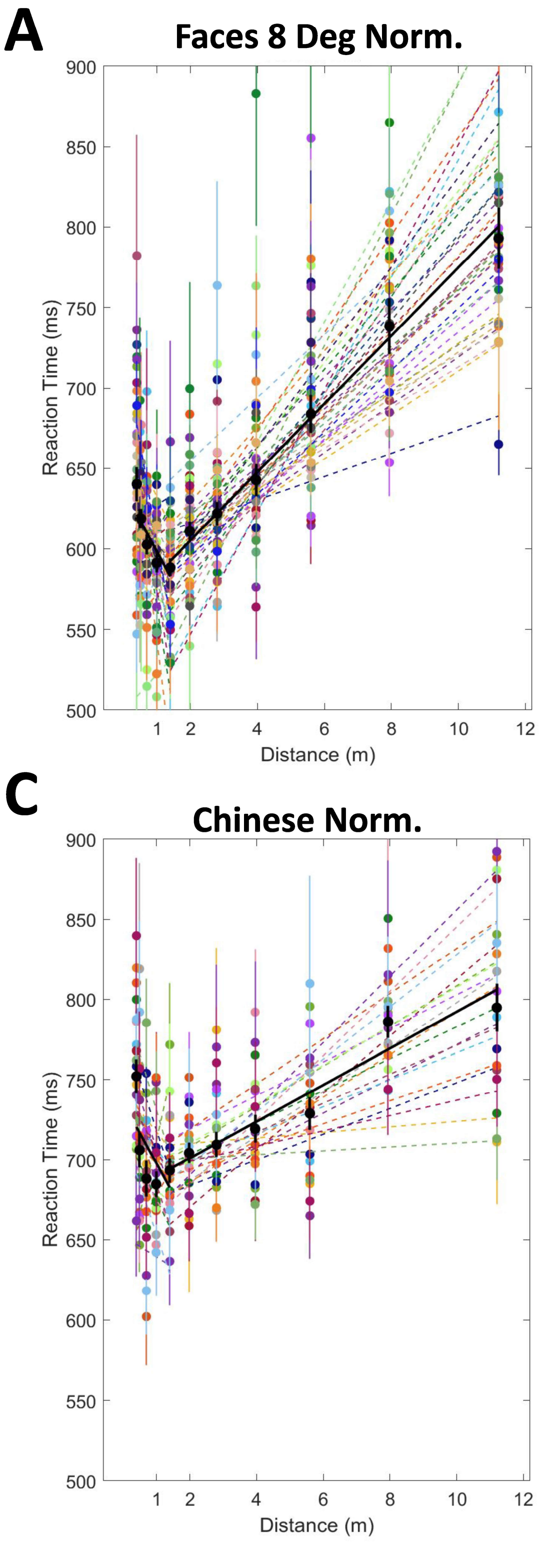

**Figure A4.** Distance conversion data for every participant from Groups A, B, and D combined. These plots show normalized data without affecting the slopes. (A) 8 Degree Faces data, with 36 participants total. (B) 4 Degree Faces data, with 30 participants total. (C) 8 Degree Chinese data, 19 participants total. (D) 8 Degree English data, 23 participants total.
